# ukbtools: An R package to manage and query UK Biobank data

**DOI:** 10.1101/158113

**Authors:** Ken B. Hanscombe, Jonathan R.I Coleman, Matthew Traylor, Cathryn M. Lewis

## Abstract

**Summary:** The UK Biobank is a resource that includes detailed health-related data on about 500,000 individuals and is available to the research community. ukbtools removes all the upfront data wrangling required to get a single dataset for statistical analysis, and provides tools to assist in quality control, query of disease diagnoses, and retrieval of genetic metadata.

**Availability:** The package is available for installation from the Comprehensive R Archive Network (CRAN), and includes a vignette describing the use of all functionality.

**Contact:**, https://github.com/kenhanscombe/ukbtools

## Introduction

The UK Biobank (UKB) is a research study of over 500,000 individuals from across the United Kingdom. Participants aged 40 to 69 were invited to one of 22 assessment centres between 2006 and 2010. A wide variety of health-related data has been collected at baseline and follow up, including cognitive function, imaging, disease diagnoses, and genome-wide genotyping data. The resource is open to applications from health scientists (http://www.ukbiobank.ac.uk/register-apply/). An approved application grants access to the UKB data, however, several obstacles limit immediate analysis of the data.

After downloading and decrypting UKB data with the supplied UKB programs, multiple files need to be integrated into a single dataset for analysis. These data files vary in format, may be very large, and have column names based on the numerical field codes of the UKB data showcase, requiring cross-referencing with an associated html file to be interpretable. ukbtools is an R package (R Core Team, 2016) that simplifies manipulation of these data files. In a single step it processes the multiple UKB files, clears the R workspace, and creates a ready-to-use dataset with meaningful column names. The package also includes tools to visualise primary demographic data for quality control (QC) purposes, query disease diagnoses, and retrieve genetic metadata for genetic association analyses.

## 1. Constructing a UKB dataset

All functions in the ukbtools package require a UKB dataset constructed with ukb_df. As such, the following workflow is an essential first step. Users download the data, decrypt it, and create a “UKB fileset” (.tab,.r,.html) with the supplied UKB programs. An example is included in the ukbtools package vignette “explore-ukb-data” and full details of the download and decrypt process are provided in the “Using UK Biobank Data” documentation (http://biobank.ctsu.ox.ac.uk/crystal/docs/UsingUKBData.pdf).

The function ukb_df takes two arguments, the stem of the fileset and optionally the full path if the fileset is in a different location, and returns a data.frame with meaningful column names. Column names are a contraction to snake_case of the full-length variable descriptions in the .html file with all punctuation and R special characters removed.

~~~
install.packages("ukbtools")
library(ukbtools)

my_ukb_data <- ukb_df("ukbxxxx", path = "/full/path/to/fileset/")
~~~

For a fileset with a 2.6 GB .tab file, processing takes approximately 10 minutes (MacBook Pro, 2.4 GHz Processor, 8 GB 1600 MHz DDR3 Memory). The rate-limiting step is R reading and parsing the code in the UKB-generated .r file to rename factor levels.

## 2. Primary demographic data

Typically, researchers will focus on a subset of the data, e.g., those individuals meeting some inclusion criteria, or with data available on a particular variable. Visualizing the demographic data for this subset of the UKB can act as a fast QC tool. ukb_context generates a single figure summary of distribution of primary demographic data for a subset of individuals relative to a reference set (Figure 1). One use for this tool is to establish representativeness. For example, comparing the distribution of the primary demographics of the subset of individuals who have data for a variable of interest to those without data (NA, or missing) for that particular variable. ukb_context also allows you to flexibly specify the comparison groups as a logical vector, e.g., body mass index greater than 40 and age less than 50 = TRUE (Subset), otherwise FALSE (Reference).

**Figure 1.**
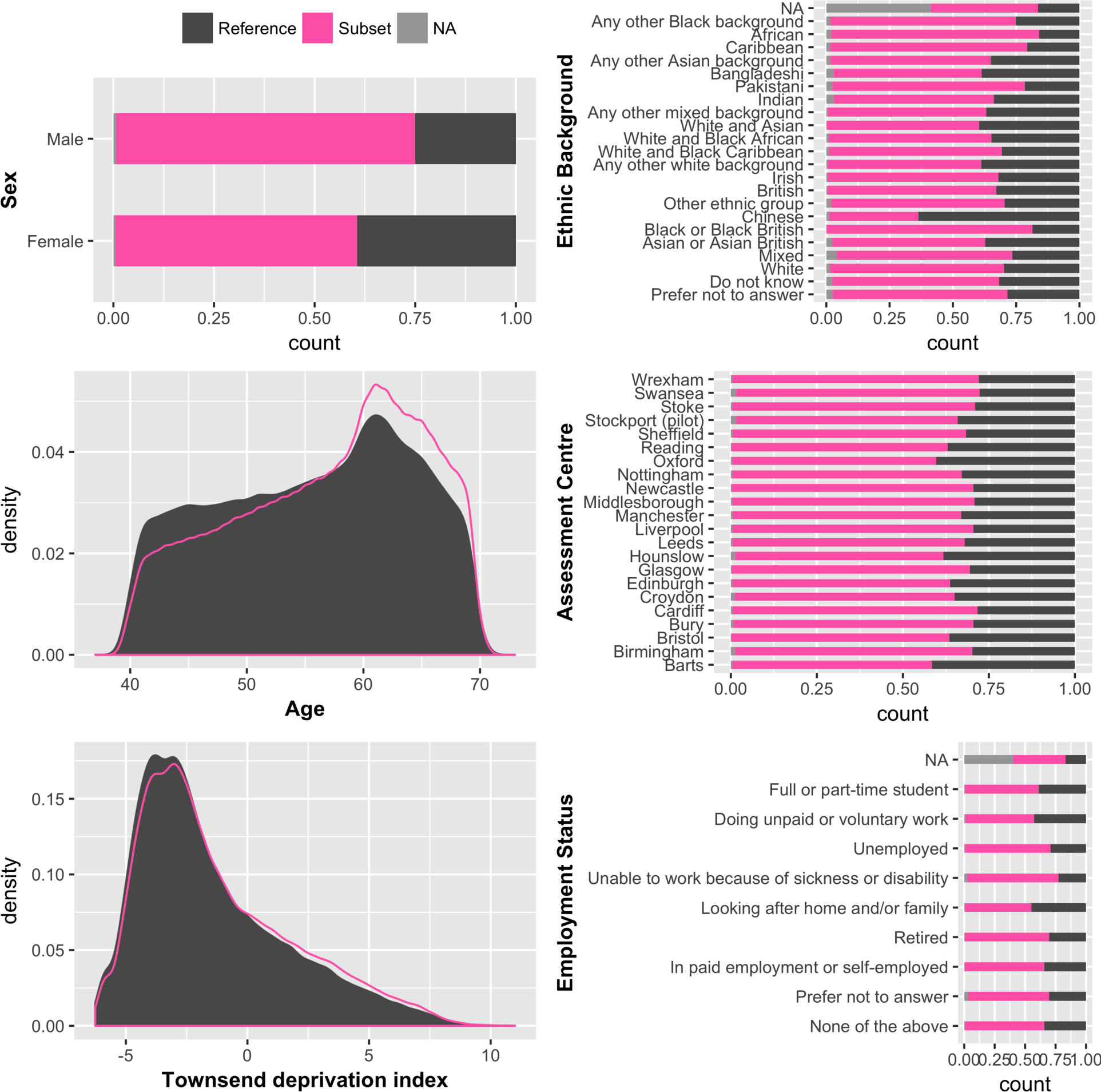
Primary demographic data for a UKB subset of interest. The subset is individuals with BMI >= 25; the reference is BMI < 25. Barplots are displayed as proportions, e.g., about 1/3 of all participants who identified as “Chinese” were overweight compared to about 2/3 of all participants who identified as “British”. ukb_context also allows the user to draw barplots as “stacked” or “side-by-side” bars representing counts, which would reveal there were many more “British” participants (442,698) than there were “Chinese” (1,574).

## 3. Disease diagnoses

UKB includes data from hospital episode inpatient statistics, useful for making disease diagnoses. ukbtooLs includes four diagnosis reference datasets to enable the interpretation of these codes: the International Statistical Classification of Diseases and Related Health Problems (ICD) revision 9 and revision 10 chapters (icd9chapters, icd10chapters) and the ICD-9 and ICD-10 codes (icd9codes, icd10codes). Respectively, these provide high-level “disease block” information (e.g. chapter 9, disease block 100-199, Diseases of the circulatory system) and granular diagnosis-specific information (e.g. code I74, Arterial embolism and thrombosis).

Two convenience functions allow the user to query both the codes and the descriptions in the included datasets. Given a particular disease code, ukb_icd_code_meaning retrieves its full description. ukb_icd_keyword returns all diseases whose descriptions include a particular keyword (actually a regular expression, e.g. “cardio”). The diagnoses of an individual can be retrieved with ukb_icd_diagnosis, which can be used as a QC tool to assess outlying individuals in an analysis. A useful exploratory analysis tool is ukb_icd_prevaLence, which returns the frequency of an ICD diagnosis code in the UKB dataset. It is possible to explore disease frequency in sub-groups of interest.

**Figure 1. Primary demographic data for a UKB subset of interest.** The subset is individuals with BMI >= 25; the reference is BMI < 25. Barplots are displayed as proportions, e.g., about 1/3 of all participants who identified as “Chinese” were overweight compared to about 2/3 of all participants who identified as “British”. ukb_context also allows the user to draw barplots as “stacked” or “side-by-side” bars representing counts, which would reveal there were many more “British” participants (442,698) than there were “Chinese” (1,574).

## 4. Genetic metadata

Effective use of the UKB genetic data requires the genetic metadata included in the UKB fileset. ukb_gen_meta retrieves assessment centre, genetic ethnicity, genetic sex, percentage SNP missingness, recommended exclusions, relatedness indicator, Affymetrix and genotyping quality control indicators variables, and genotyping chip. Assessment centre is numerically coded. ukb_gen_meta returns a dataset with the original coding and a variable with centre names. ukb_gen_centre is also provided as a standalone function to add centre names to any dataset.

A list of IDs for recommended exclusions and heterozygosity outliers (+/- 3*SD) can be retrieved with ukb_gen_excl and ukb_gen_het respectively. Setting the parameter all.het = TRUE causes ukb_gen_het to return heterogeneity statistics for all samples. ukb_gen_pcs retrieves the twenty principal components used to control for population structure in the genetic data. ukb_gen_rel returns a data.frame with id, pair (a numeric identifier for related pairs), and kinship (kinship coefficient). Users can create a table of counts for different degrees of relatedness (e.g. monozygotic twins, full siblings), and reproduce the relatedness plot on page 15 of the UKB documentation (http://www.ukbiobank.ac.uk/wp-content/uploads/2014/04/UKBiobank_genotyping_QC_documentation-web.pdf) for any subset of the full UKB data with ukb_gen_rel_count.

Ukbtools takes all the time-consuming effort out of preparing input data for PLINK (Chang C.C., 2015, http://www.cog-genomics.org/plink/1.9/) and BGENIE (Bycroft et al., 2017, https://jmarchini.org/bgenie/) analyses with a set of read (fam and sample files) and write (phenotype, covariate and exclusions files) functions.

## Conclusion

Having a dataset with meaningful variable names, a set of UKB-specific exploratory data analysis tools, and a set of helper functions to retrieve and write genetic metadata to file, will rapidly enable UKB users to undertake their research.

## Acknowledgement

This research has been conducted using the UK Biobank Resource (Application 13427).

